# Emergence of new subgenomic mRNAs in SARS-CoV-2

**DOI:** 10.1101/2022.04.20.488895

**Authors:** Harriet V Mears, George R Young, Theo Sanderson, Ruth Harvey, Margaret Crawford, Daniel M Snell, Ashley S Fowler, Saira Hussain, Jérôme Nicod, Thomas P Peacock, Edward Emmott, Katja Finsterbusch, Jakub Luptak, Emma Wall, Bryan Williams, Sonia Gandhi, Charles Swanton, David LV Bauer

**Affiliations:** RNA Virus Replication Laboratory, The Francis Crick Institute, London, UK; Bioinformatics and Biostatistics STP, The Francis Crick Institute, London, UK; Malaria Biochemistry Laboratory, The Francis Crick Institute, London, UK; Worldwide Influenza Centre, The Francis Crick Institute, London, UK; Advanced Sequencing Facility, The Francis Crick Institute, London, UK; Department of Infectious Disease, St Mary’s Hospital, Imperial College London, London, UK; Centre for Proteome Research, Department of Biochemistry & Systems Biology, Institute of Systems Molecular & Integrative Biology, University of Liverpool, Liverpool, UK; Immunoregulation Laboratory, The Francis Crick Institute, London, UK; MRC Laboratory of Molecular Biology, Cambridge, UK; Crick/UCLH Legacy Study, The Francis Crick Institute, London, UK; University College London and National Institute for Health Research (NIHR) University College London Hospitals (UCLH) Biomedical Research Centre, London, UK; Neurodegeneration Biology Laboratory, The Francis Crick Institute, London, UK; Cancer Evolution and Genome Instability Laboratory, The Francis Crick Institute, London, UK; Genotype-to-Phenotype (G2P-UK) National Virology Consortium

## Abstract

Two mutations occurred in SARS-CoV-2 early during the COVID-19 pandemic that have come to define circulating virus lineages^1^: first a change in the spike protein (D614G) that defines the B.1 lineage and second, a double substitution in the nucleocapsid protein (R203K, G204R) that defines the B.1.1 lineage, which has subsequently given rise to three Variants of Concern: Alpha, Gamma and Omicron. While the latter mutations appear unremarkable at the protein level, there are dramatic implications at the nucleotide level: the GGG→AAC substitution generates a new Transcription Regulatory Sequence (TRS) motif, driving SARS-CoV-2 to express a novel subgenomic mRNA (sgmRNA) encoding a truncated C-terminal portion of nucleocapsid (N.iORF3), which is an inhibitor of type I interferon production. We find that N.iORF3 also emerged independently within the Iota variant, and further show that additional TRS motifs have convergently evolved to express novel sgmRNAs; notably upstream of Spike within the nsp16 coding region of ORF1b, which is expressed during human infection. Our findings demonstrate that SARS-CoV-2 is undergoing evolutionary changes at the functional RNA level in addition to the amino acid level, reminiscent of eukaryotic evolution. Greater attention to this aspect in the assessment of emerging strains of SARS-CoV-2 is warranted.

---

SARS-CoV-2 has continued to evolve since its emergence in the human population^1^. Two notable mutations occurred early in the pandemic that define circulating lineages: first, a change in Spike (D614G) that is now present in 99% of all sequences and defines the B.1 lineage in conjunction with the polymerase mutation nsp12:P323L; and second, a double substitution in nucleocapsid (R203K,G204R) that is present in a third of all sequences and is the defining mutation of viruses in the B.1.1 lineage and its descendants — including three Variants of Concern: Alpha (B.1.1.7), Gamma (P.1), and Omicron (B.1.1.529) (**Fig. 1a**).

**Fig. 1.**
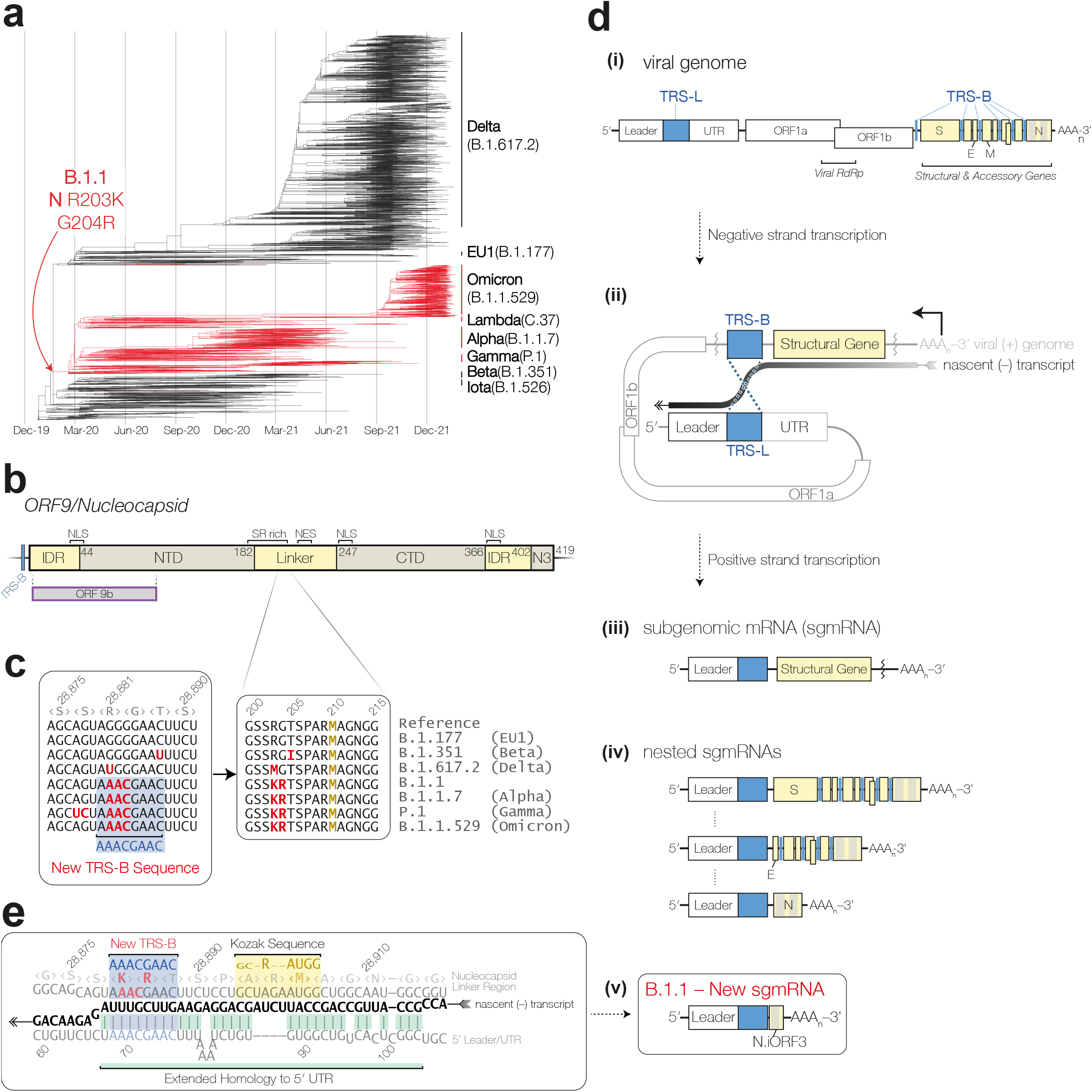
Nucleocapsid R203K, G204R mutations in the SARS-CoV-2 B.1.1 lineage generate a novel TRS-B site and new subgenomic mRNA (sgmRNA). **a**, Phylogenetic reconstruction of SARS-CoV-2 evolution in humans, with lineage-defining mutations for B.1 and B.1.1 indicated. Viruses with Nucleocapsid K204R mutation (B.1.1 and its descendants) are coloured in red. **b**, Diagram of nucleocapsid protein domains. The N-terminal (NTD), linker, C-terminal (CTD), and N3 domains are shown, along with intrinsically disordered regions (IDR), serine-arginine (SR) rich region. Reported nuclear localisation signals (NLS), nuclear export signal (NES), the extent of overlapping open reading frame 9b (ORF9b), and upstream transcription regulatory sequence (TRS-B) are also indicated. **c**, Sequence alignment of amino acids 200-215 of nucleocapsid, and alignment of the corresponding nucleotide sequences show emergence of a new TRS-B motif. **d**, Coronavirus structural (S, E, M, N) and accessory genes (yellow highlight) are expressed by discontinuous transcription during (–) strand synthesis, by template switching at TRS-B sequences (blue highlight), upstream of the structural or accessory genes, to the homologous TRS-L in the 5′UTR. Since the efficiency of template switching is not 100%, this process generates a set of nested subgenomic RNAs. In the B.1.1 lineage, the new TRS-B leads to an additional sgmRNA expressing N.iORF3, a C-terminal fragment of nucleocapsid. **e**, The sequence context of the novel N.iORF3 sgmRNA, showing TRS-B (blue highlight), extended homology to the 5′UTR (green highlight) during nascent (–) strand RNA synthesis (black), and downstream start codon and Kozak context (yellow highlight). The phylogenetic tree in panel a was adapted from Nextstrain^32,33^.

The coronavirus nucleocapsid is a multifunctional protein with a highly conserved domain architecture^2^ (**Fig. 1b**): it consists of two structured domains (Nterminal domain, NTD; and C-terminal domain, CTD) separated by a disordered serine-arginine (SR)-rich linker and flanked by two intrinsicallydisordered domains. The NTD, SR-rich linker, and CTD bind to RNA, and the extreme C-terminus also contains a short domain (“N3”) that interacts with the viral membrane protein^3^ and is thought to facilitate viral genome packaging by anchoring the viral RNA to nascent virions^4^. The R203K,G204R mutations appear unremarkable on an amino acid level, but have important implications at the nucleotide level: the underlying substitutions (G28881A, G28882A, G28883C) create a new transcription regulatory sequence^5^ (A<u>GGG</u>GAAC→A<u>AAC</u>GAAC, **Fig. 1c**).

Transcription regulatory sequences (TRSes) are well-characterised cis-acting RNA regulatory sequences, which are required for expression of the structural (spike, envelope, membrane, nucleocapsid) and accessory genes in the 3′ end of coronavirus genomes, downstream of ORF1ab^6^ (**Fig. 1d.i**). During negative strand transcription, as the viral polymerase copies a TRS in the body of the genome (TRS-B), the nascent RNA strand can dissociate from the genomic template and re-anneal to the homologous TRS in the 5′ leader (TRS-L), whereupon transcription is re-initiated (**Fig. 1d.ii**). The resultant antisense subgenomic RNAs are then copied back to positive sense, yielding a nested set of subgenomic messenger RNAs (sgmRNAs) with a 5′ Leader, TRS, the relevant ORF, and the remaining 3′ portion of the genome (**Fig. 1d.iii**), which are 5′ and 3′ co-terminal with one-another (**Fig. 1d.iv**). This mechanism of discontinuous transcription, which is found throughout the order *Nidovirales*, allows the virus to tune sgmRNA expression levels by increasing or decreasing the degree of homology between the regions flanking the TRS-L and the TRS-B — or, more precisely, the strength of base-pairing between the TRS-L and the anti-TRS-B in the nascent negative strand RNA^7^.

The new TRS-B in the B.1.1 lineage is flanked by substantial sequence homology to the 5′ Leader (4 nt directly adjacent to the TRS, plus distal structures, **Fig. 1e**). The resultant sgmRNA (**Fig. 1d.v**) contains a start codon, in good Kozak sequence context for translation initiation, in frame with the nucleocapsid open reading frame at Met-210 (**Fig. 1e**). This creates a new open reading frame that encodes residues 210-419 of nucleocapsid, fully encompassing the CTD and C-terminal N3 domain. Since this is the third internal open reading frame identified within the nucleocapsid coding region of SARS-CoV-2, we have named it N.iORF3 (“Nigh ORF Three”, rhymes with English “sigh”), following the naming system used by Finkel *et al*.^8^

To verify whether the N.iORF3 sgmRNA is transcribed during infection in cell culture, we infected Vero E6 cells at a high multiplicity with either a very early 2020 UK isolate (B lineage), a late 2020 UK isolate (Alpha variant, B.1.1.7 lineage), a late 2020 South African isolate (Beta variant, B.1.351 lineage), or a late 2021 UK isolate (Omicron BA.1 variant, B.1.1.529.1 lineage) and harvested cells at 24 hours post-infection. RNA was extracted and analysed by RT-PCR using a forward primer against the 5′ Leader and a reverse primer against the 3′-end of N (**Fig. 2a**). When analysed by agarose gel electrophoresis, products corresponding to the canonical full-length N sgmRNA were detected in all SARS-CoV-2 infected cells, and an additional shorter product corresponding to the N.iORF3 sgmRNA was detected in Alpha- and Omicroninfected cells, but not lineage B-, and Beta-infected cells (**Fig. 2b**).

**Fig. 2.**
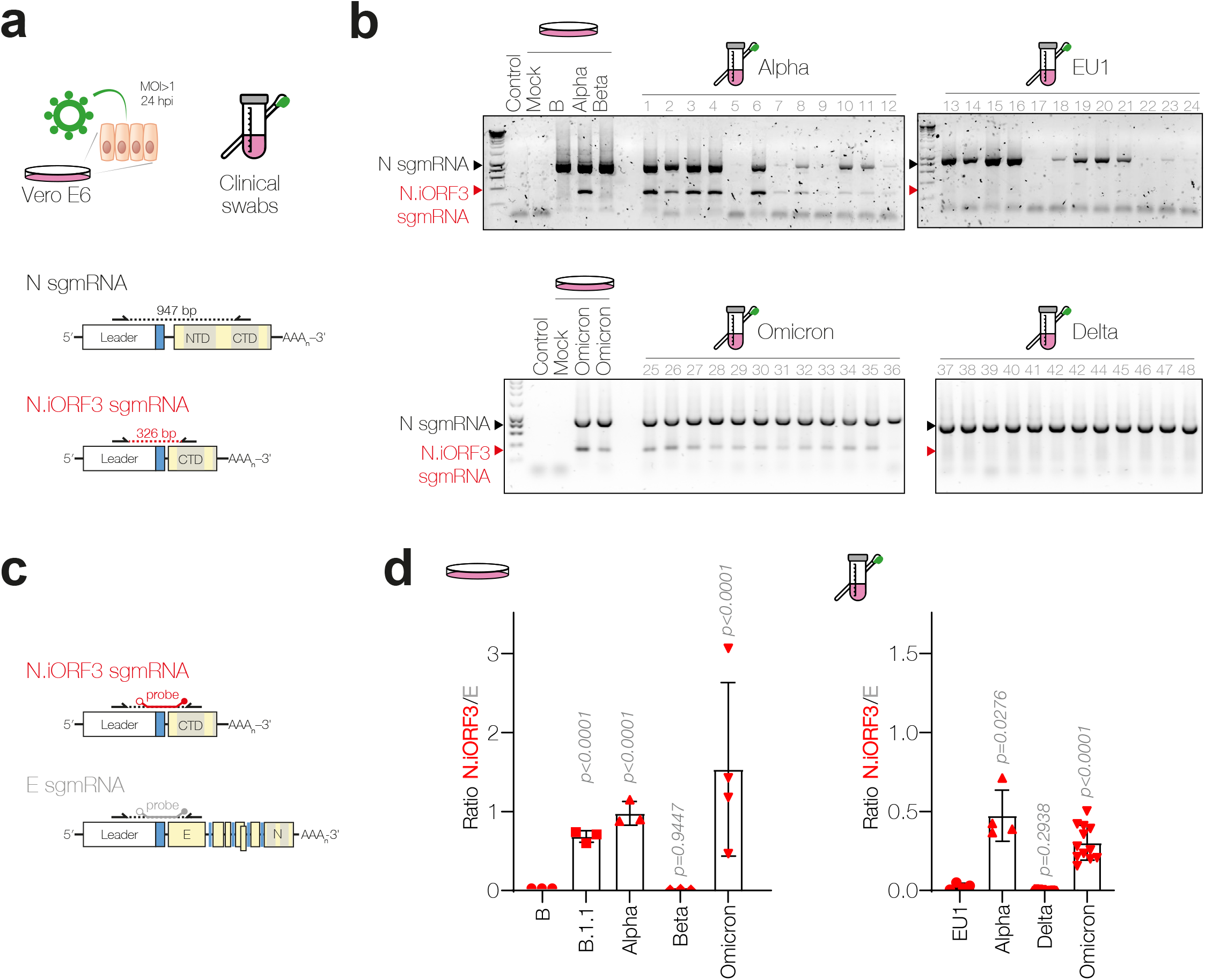
N.iORF3 sgmRNA is produced during SARS-CoV-2 infections in cell culture and in humans. **a**, Schematic representation of RT-PCR analysis of RNA extracted from infected VeroE6 cells at 24 hours post infection or nasopharyngeal swabs, showing positions of primers. **b**, RT-PCR detection of canonical nucleocapsid (N) and N.iORF3 sgmRNAs in RNA extracted from infected cells and clinical swabs, for SARS-CoV-2 variants as indicated. **c**, Schematic representation of qRT-PCR primer probe sets for N.iORF3 and envelope (E) sgmRNAs. **d**, RT-qPCR analysis of N.iORF3 sgmRNA copy number, expressed as a ratio of E copy number in infected cells and human clinical swabs. For infected VeroE6 cells, data are means and standard deviations of at least three biological replicates, compared to ‘Lineage B’ or to Alpha by one-way ANOVA and Dunnett’s test. For clinical swabs, data are means and standard deviations of four (EU1/Alpha) or twelve (Delta/Omicron) swab samples per lineage, compared by one-way Brown-Forsythe and Welch ANOVA and Dunnett’s T3 test, to account for unequal variances. P values are shown.

We then sought to confirm the presence of the N.iORF3 sgmRNA in human infections. We analysed occupational health screening swab samples from UK National Health Service (NHS) healthcare workers and employees of the Francis Crick Institute processed by the Crick COVID-19 Consortium Testing Centre and retained for analysis as part of the Legacy study (Cohort A1, NCT04750356). The Legacy study was approved by London Camden and Kings Cross Health Research Authority Research and Ethics committee (IRAS number 286469) and is sponsored by University College London Hospitals. We analysed 12 samples from each of the main COVID-19 “waves” from late 2020 to early 2022: B.1.177 (EU1, autumn 2020); B.1.1.7 (Alpha, winter 2020/1); B.1.617.2 (Delta, summer 2021); and BA.1 (Omicron, winter 2021/2). Analysis by RT-PCR revealed that N.iORF3 sgmRNA was specifically present in human clinical samples from the B.1.1 lineage (Alpha, Omicron) but not from the B.1 lineage (EU1, Delta) (**Fig. 2b**).

To quantify the amount of N.iORF3 sgmRNA present in samples and to avoid quantitative biases from end-point PCR^9^, we designed an RT-qPCR assay to compare N.iORF3 sgmRNA with other viral RNA species. Probes spanned the leader-TRS-sgmRNA junction (**Fig. 2c**), allowing absolute quantitation of N.iORF3, N, and Envelope (E) sgmRNA copy number (**Extended Data Fig. 1a-c**). N.iORF3 sgmRNA was consistently expressed at an equivalent level (30-150%) to E sgmRNA (**Fig. 2d**), approximately 100-fold lower than the highly abundant N sgmRNA, in both clinical swabs and infected cells (**Extended Data Fig. 1d-e**). A previous report^10^ suggested that the Alpha variant also expressed a distinct ORF9b-specific sgmRNA. While we were able to detect this product by nanopore sequencing of endpoint PCR products, its level of expression was below the limit of detection (10^2^ copies) of our RT-qPCR assay using ORF9bspecific sgmRNA probes (**Extended Data Fig. 2**).

We next sought to determine whether a protein product was produced from N.iORF3 (**Fig. 3a**) in infected cells. VeroE6 ACE2-TMPRSS2 cells were infected with Lineage B, B.1.1, Alpha, Beta or Delta viruses, and harvested at 20 hours post infection. When analysed by immunoblotting, N expression was largely consistent between variants and we also observed a band at ∼25 kDa in B.1.1 and Alpha-infected cells (**Fig. 3b**), consistent with the mobility of recombinant N.iORF3 protein (**Fig. 3c**). In SARS-CoV-1, type I interferon production is strongly inhibited by N, specifically its C-terminal domain (CTD), which sequesters double-stranded RNA in the cytoplasm away from host pattern recognition receptors^11,12^ (**Fig. 3a**). We therefore hypothesised that N.iORF3 protein, which encompasses the CTD of N, might act as an interferon antagonist. To test this hypothesis, HEK293T cells were transfected with plasmids encoding N or N.iORF3, or influenza A virus NS1 protein as a positive control; after 24 hours, cells were transfected with poly(I:C), a synthetic double-stranded RNA analogue which stimulates innate immune signalling. Cells transfected with either N or N.iORF3 expressed lower levels of IFNb and IFIT1 mRNA (**Fig. 3d**), indicating that N.iORF3 can antagonise IFN signalling downstream of dsRNA sensing in the cytoplasm.

**Fig. 3.**
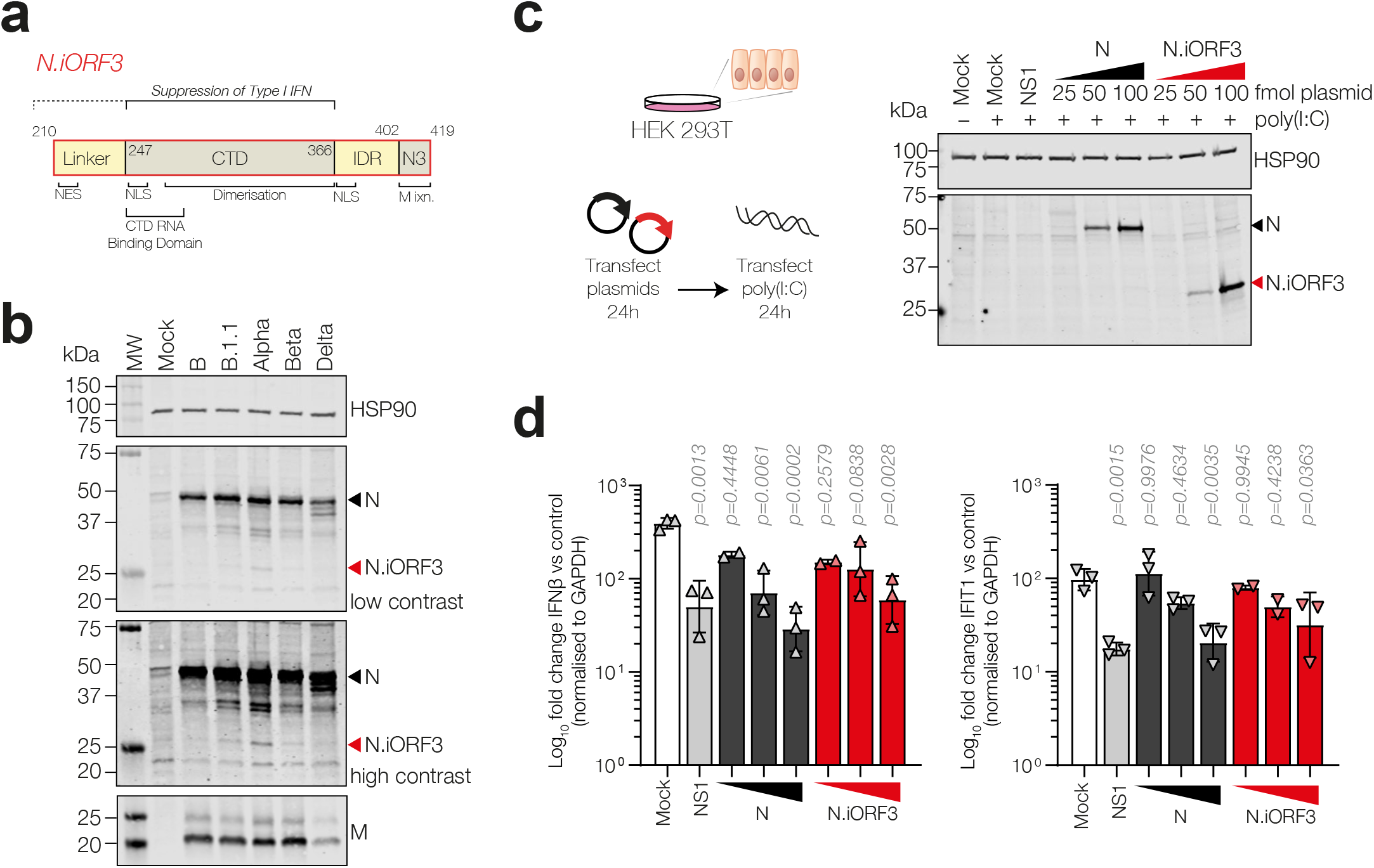
N.iORF3 sgmRNA expresses N.iORF3 protein, a truncated form of nucleocapsid which acts as an innate immune signalling antagonist. **a**, Diagram of N.iORF3 protein, showing domains from nucleocapsid and described functions. “M ixn.” indicates region of interaction with the viral membrane protein. **b**, Western blot analysis of lysates from infected VeroE6 ACE2-TMPRSS2 cells. N.iORF3 is indicated with red arrowheads. **c**, Schematic representation of immune interference assay and expression of N and N.iORF3 in transfected HEK293T cells. **d**, Expression of type I interferon (IFNb, left panel) and a representative interferon-stimulated gene (IFIT1, right panel), normalised to housekeeping gene expression (GAPDH) and expressed as fold change in cells transfected with poly(I:C) compared to untransfected cells, in the presence of increasing concentrations of N-or N.iORF3-expressing plasmids (25, 50 or 100 fmol) or NS1 from influenza A virus as a positive control (100 fmol). Data are means and standard deviations of at least two biological replicates, compared to mock-transfected cells by one-way ANOVA and Dunnett’s test. P values are shown.

We wondered whether N.iORF3 sgmRNA might have evolved independently outside of the B.1.1 lineage, and therefore used Taxonium (see Methods) to search public SARS-CoV-2 genomes within the complete UShER phylogenetic tree^13^ that contained the R203K,G204R mutation in nucleocapsid, but lay outside of the B.1.1 lineage. We found examples of convergent evolution of the N.iORF3 sgmRNA, notably within the Iota variant (B.1.526, **Extended Data Fig. 3a**), where samples clustered within geographic regions, showed evidence of ongoing transmission, and were detected by multiple depositing laboratories (**Extended Data Table 1**). Intriguingly, we also observed multiple independent instances of further evolution of the N.iORF3 TRS region (a silent double-nucleotide substitution at S202) that increases homology to the 5′UTR (**Extended Data Fig. 3b**), notably in the entire Gamma lineage, at least 6 times within the Alpha lineage, and, most recently, within Omicron (**Extended Data Fig. 3c** and **Extended Data Table 1**).

We then considered whether novel TRS-B emergence might occur more widely in the SARS-CoV-2 genome, and searched deposited sequences for novel TRS-B sites based on the emergence of the AAACGAAC motif (**Fig. 4a**). Surprisingly, we found that after N.iORF3, the most frequent site is located at the end of ORF1ab, within the nsp16 coding region, 251 nt upstream of Spike.

**Fig. 4.**
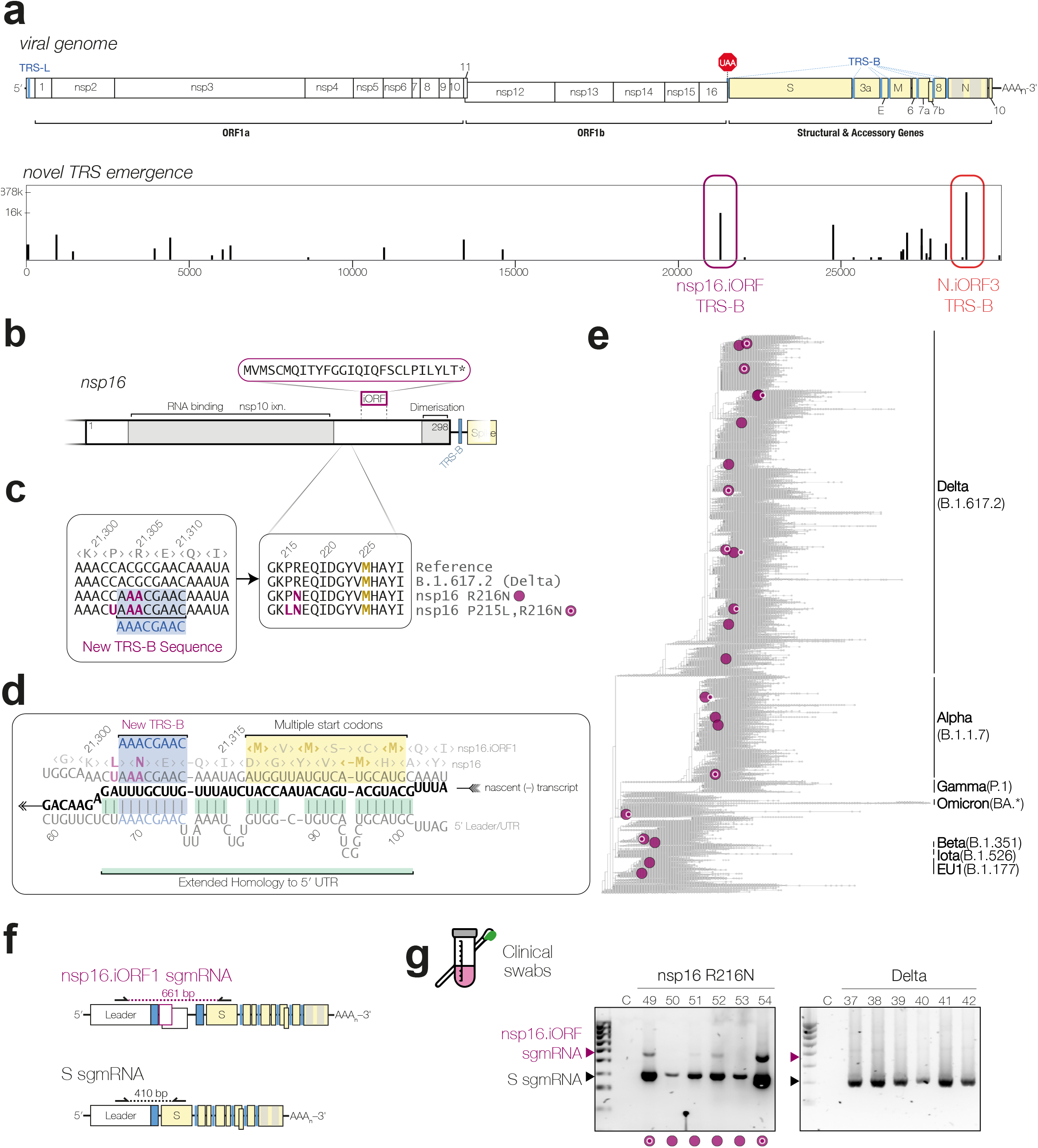
Novel TRS-B sites continue to emerge during the SARS-CoV-2 pandemic. **a**, Schematic of the SARS-CoV-2 genome (upper panel) and frequency of emergence of the TRS-B sequence (AAACGAAC) in the global SARS-CoV-2 population (lower panel). **b**, Diagram of the nsp16 coding region, including a potential transframe product. **c**, Sequence alignment of amino acids 213-228 of nsp16, and alignment of the corresponding nucleotide sequences show emergence of a new TRS-B sequence. **d**, The sequence context of the novel nsp16.iORF sgmRNA, showing TRS-B (blue highlight), extended homology to the 5′UTR (green highlight) during nascent (–) strand RNA synthesis (black), and downstream start codon and Kozak context (yellow highlight). **e**, Phylogenetic reconstruction of SARS-CoV-2 evolution in humans, with independent emergences of nsp16.iORF TRS sequence with ≥100 descendant genomes highlighted in purple (see **Extended Data Table 1**). Emergence of extended homology to the 5′UTR is indicated with white outline “bullseye” pattern. f, Schematic representation of RT-PCR analysis of RNA extracted from nasopharyngeal swabs, showing positions of primers. g, RT-PCR detection of nsp16.iORF sgmRNA in clinical swabs, indicated by purple arrowheads, or Spike sgmRNA, indicated by black arrowheads.

nsp16 is a 2′-O-methyltransferase involved in viral RNA capping, and consists of a single globular domain which binds to RNA and the methyl donor Sadenosyl methionine, as well as its cofactor, nsp10 (**Fig. 4b**)^14,15^. These functions are primarily mediated by the N-terminal two-thirds of the protein, while the C-terminal tail has a role in nsp16 dimerisation by forming a domain-swapped beta sheet with a second nsp16 subunit. A two-nucleotide substitution underlying amino acid mutation nsp16:R216N (ORF1ab:R7014N) creates a consensus TRS-B (**Fig. 4c**), immediately upstream of four tandem start codons (**Fig. 4d**). The first, second and fourth start codons are in the +1 reading frame, in moderately good Kozak sequence context, and encode a short transframe peptide, that we designated nsp16.iORF1; the third start codon, also in moderate Kozak context, is in frame with the main nsp16 ORF and encodes a C-terminal portion of nsp16, designated nsp16.iORF2.

We found at least 21 occurrences of convergent evolution of this nsp16.iORF TRS-B (**Fig. 4e** and **Extended Data Table 1**), notably including the summer 2020 B.1.1.44 outbreak in Scotland (82% of sequences), a large Alpha subclade focused on the Canadian Prairies (subsequently displaced by Delta) and a large AY.4.2 Delta subclade focused on England (subsequently displaced by Omicron). As with the N.iORF3 TRS, we also observed further evolution of the sequence surrounding the nsp16.iORF TRS that increases base-pairing to the 5′UTR, results in a further mutation at nsp16:P215L, and is present in 48% of sequenced genomes with the nsp16.iORF TRS.

To determine whether the nsp16.iORF sgmRNA is expressed in infection, we searched archived positive clinical swabs from the Legacy study that had been sequenced, and found six with the nsp16 R216N mutation, collected September 2020 – January 2021. We analysed RNA extracted from these swabs by RT-PCR using primers in the 5′ leader and in the 5′-portion of Spike coding region (**Fig. 4f**). We observed amplification of the canonical longer nsp16.iORF-specific product in four samples, while the remaining two were below the limit of detection (**Fig. 4g**). These results confirmed that the nsp16.iORF sgmRNA is expressed during human infection.

It worth noting that the repertoire of TRS sites may be larger than that considered so far: the third-most common novel TRS has evolved independently at least 7 times via a single nucleotide change within the connector domain^16^ of Spike/S2 (**Extended Data Fig. 4** and **Extended Data Table 1**), and it also contains extended homology to the 5′UTR and lies upstream of tandem start codons of a transframe peptide. Subgenomic RNAs may also be produced with non-consensus TRSes^17^; the canonical E sgmRNA, for example, is generated via a minimal ACGAAC sequence. This shorter core TRS has evolved 13 times at two distinct loci in SARS-CoV-2 upstream of E, within the disordered CTD of ORF3a, and has come to predominate in the large AY.31.1 Delta lineage, accounting for over 30% of all Delta sequences in Australia (**Extended Data Fig. 5** and **Extended Data Table 1**). Together, our findings imply that TRS site emergence within SARS-CoV-2 happens frequently on a global scale can generally lead to novel sgmRNA expression.

The emergence of novel open reading frames — and concomitant TRS-B evolution — is a hallmark of coronavirus evolution and adaptation to new hosts^18–20^. This evolution may occur via horizontal gene transfer between coronaviruses, such as mobile modules within the Spike gene implicated in expansion of viral host range, and via nonhomologous recombination with unrelated viruses, such as the acquisition of a haemagglutinin esterase gene by an ancestral betacoronavirus from an influenza C-like virus. Coronaviruses may also duplicate parts of their own genomes: in SARS-CoV-2 and related viruses, ORF3a diverged from a copy of the membrane gene^21^, while in mouse coronavirus, functional redundancy between the membrane and envelope genes suggests a common evolutionary origin^22^, with each evolving distinct TRS-B sequences. Moreover, the 3′ regions of ORF6 and ORF8 in SARS-CoV-2, have significant sequence homology to the 5′UTR, while in HCoV-OC43 the intergenic region between Spike and the ns12.9 genes has 73% nucleotide identity to the 5′ end, suggesting that the TRS itself and flanking sequence can serve as a source of new genetic material. These processes — reminiscent of eukaryotic gene evolution via exon shuffling and duplication of promoter and regulatory elements^23,24^ — are fundamental to evolution in nidoviruses^25,26^ and RNA viruses more broadly^27^.

In the case of the novel sgmRNAs identified here in SARS-CoV-2, the complete set of functions of the proteins and peptides they encode is not yet known. Both nsp16.iORF2 and N.iORF3 may be regulated differently in cells, as they lack domains of their fulllength protein counterparts. Conversely, they may also regulate full-length protein function since both contain the dimerization domains of their full-length cognates, raising the possibility of heterodimers of unknown function or competition for the dimerization domain of full-length protein. The coding changes introduced by the TRS site itself may also affect the function of the parental protein. For example, the region of N in sarbecoviruses surrounding the N.iORF3 TRS has been variously implicated in linker phosphorylation^28^, genome packaging^29^, and host cell cycle arrest^30^. Perhaps most fundamentally, the emergence of novel TRSes may act on an RNA level — just as the introduction of alternate splice sites may act during eukaryotic transcription — to tune the overall programme of gene expression, which is still being refined in SARS-CoV-2 as new variants emerge^31^.

Together, our results highlight an important feature of SARS-CoV-2 evolution at the functional RNA level, with consequences for understanding both the past and future of the COVID-19 pandemic. The convergent evolution of TRSes makes it more challenging to trace the early evolution of SARS-CoV-2 in humans in which the N:203K,204R mutation was often used as a lineage-defining mutation. Going forwards, it will be important to ensure that genomic surveillance pipelines and tools such as Nextstrain^32,33^, GISAID^34^, and CoV-GLUE^35^ will allow easy identification of functionally-important nucleotide-level changes in addition to the oft-scrutinised amino acid mutations.

## Acknowledgements

This research was funded in whole, or in part, by the Wellcome Trust [FC011104, FC011233, 210918/Z/18/Z]. For the purpose of Open Access, the author has applied a CC BY public copyright licence to any Author Accepted Manuscript version arising from this submission. The phylogenetic reconstructions depicted in Fig. 1a and Extended Data Fig. 3a-b are derivatives of those produced by nextstrain.org^32,33^, used under CC BY. We thank all researchers who have submitted SARS-CoV-2 genomes to the GISAID database: an acknowledgement table for the specific set analysed here can be found in **Extended Data Table 2**. We thank Simon Caidan, Robert Goldstone, Maria Greco, and Michael Bennett for technical assistance; and Steve Gamblin, George Kassiotis, Wendy Barclay, and Ervin Fodor for helpful discussions. We thank Wendy Barclay, ‘Assessment of Transmission and Contagiousness of COVID-19 in Contacts’ (ATACCC), the NIHR Health Protection Research Unit in Respiratory Infections, Imperial College London (NIHR200927), Public Health England, Leo James, Steve Goodbourne, Tulio de Oliveira, Alex Sigal, Khadija Kahn, Thushan de Silva, Gavin Screaton, and the G2P-UK National Virology Consortium for support and rapid sharing of reagents and samples, as well as the participants of the Crick SARS-CoV-2 Longitudinal Study: Understanding Susceptibility, Transmission and Disease Severity (Legacy Study). The Legacy Study is supported by the NIHR University College London Hospitals Biomedical Research Centre. Theo Sanderson is supported by grant 210918/Z/18/Z from the Wellcome Trust. Jakub Luptak is supported by UK Medical Research Council and UK Research and Innovation. This work was supported by the Francis Crick Institute which receives its core funding from Cancer Research UK (FC011104, FC011233), the UK Medical Research Council (FC011104, FC011233), and the Wellcome Trust (FC011104, FC011233).

## Competing Interests

The authors declare no competing interests directly related to this work. CSw reports interests unrelated to this work: grants from BMS, Ono-Pharmaceuticals, Boehringer-Ingelheim, Roche-Ventana, Pfizer and Archer Dx, unrelated to this Correspondence; personal fees from Genentech, Sarah Canon Research Institute, Medicxi, Bicycle Therapeutics, GRAIL, Amgen, AstraZeneca, BMS, Illumina, GlaxoSmithKline, MSD, and Roche-Ventana, unrelated to this Correspondence; and stock options from Apogen Biotech, Epic Biosciences, GRAIL, and Achilles Therapeutics, unrelated to this Correspondence. DLVB reports grants from AstraZeneca unrelated to this work. All other authors declare no competing interests.

## Methods

### Cell Lines, Viruses, and In Vitro Infections

Vero E6 (Pasteur), Vero V1 (a gift from Stephen Goodbourn) and VeroE6 ACE2-TMPRSS2 cells^36^ (a gift from Suzannah Rihn) were maintained in Dulbecco’s Modified Eagle Medium, supplemented with 10% foetal calf serum and penicillin-streptomycin (100 U/mL each). The SARS-CoV-2 B lineage isolate used (hCoV-19/England/02/2020) was obtained from the Respiratory Virus Unit, Public Health England, UK, (GISAID accession EPI_ISL_407073). The B.1.1 lineage strain used was isolated from a healthcare worker swab as part of the Legacy study and has the genotype: C241T, C3037T, nsp12: P323L, S: D614G, N: S194L, N: R203K, N: G204R. The SARS-CoV-2 B.1.1.7 isolate (“Alpha”) was hCoV-19/England/204690005/2020, which carries the D614G, Δ69-70, Δ144, N501Y, A570D, P681H, T716I, S982A and D1118H mutations in Spike, and was obtained from Public Health England (PHE), UK, through Prof. Wendy Barclay, Imperial College London, London, UK through the Genotype-to-Phenotype National Virology Consortium (G2P-UK). The B.1.617.2 (“Delta”) isolate was MS066352H (GISAID accession number EPI_ISL_1731019), which carries the T19R, K77R, G142D, Δ156-157/R158G, A222V, L452R, T478K, D614G, P681R, D950N mutations in Spike, and was kindly provided by Prof. Wendy Barclay, Imperial College London, London, UK through the Genotype-to-Phenotype National Virology Consortium (G2P-UK). The BA.1 (“Omicron”) isolate was M21021166, which carries the A67V, Δ69-70, T95I, Δ142-144, Y145D, Δ211, L212I, G339D, S371L, S373P, S375F, K417N, N440K, G446S, S477N, T478K, E484A, Q493R, G496S, Q498R, N501Y, Y505H, T547K, D614G, H655Y, N679K, P681H, A701V, N764K, D796Y, N856K, Q954H, N969K, and L981F mutations in Spike, and was kindly provided by Prof. Gavin Screaton, University of Oxford, Oxford, UK through the Genotype-to-Phenotype National Virology Consortium (G2P-UK). The B.1.351 isolate (“Beta”) was obtained from Alex Sigal and Tulio de Olivera. Viral genome sequencing of this B.1.351 identified S: Q677H and S: R682W mutations at the furin cleavage site in ∼45% of genomes. Virus stocks were propagated in Vero V1 cells by infection at an MOI of 0.01 in Dulbecco’s Modified Eagle Medium, supplemented with 1% foetal calf serum and penicillin-streptomycin (100 U/mL each), harvested when CPE was visible, and stocks were titrated on Vero E6 cells. For in vitro infections to examine sgmRNA production, Vero E6 cells were infected at an MOI >1 in Dulbecco’s Modified Eagle Medium, supplemented with 1% foetal calf serum and penicillinstreptomycin (100 U/mL each). At 7 or 24 hours postinfection, cells were washed with phosphate-buffered saline and lysed in TRIzol or Laemmli buffer. RNA from infected cells was extracted using a Direct-zol RNA MiniPrep kit (Zymo Research).

### Plasmids

Sequences for N and N.iORF3 were amplified from cDNA from B.1.1.7-infected cells, to include 5′ NheI and 3′ NotI sites, then ligated into pCDNA3-T2A-mCherry, to generate pCDNA3-B117-N-T2A-mCherry and pCDNA3-B117-Nstar-T2A-mCherry. pCDNA3-NS1 was a kind gift from Caetano Reis e Sousa.

### Clinical samples

Extracted RNA from occupational health screening swab samples of UK National Health Service (NHS) healthcare workers at the Crick COVID Consortium Testing Centre18 was obtained from the Crick/UCLH SARS-CoV-2 Longitudinal Study (Legacy Study) [COVID-19] (IRAS ID 286469). These samples were collected between December 2020 and February 2021 and had tested positive for SARS-CoV-2 (TaqPath assay, Thermo Fisher) and the viral genomes had been fully sequenced (ARTIC v333,34, GridION), and lineage assigned using pangolin^1^. The 12 samples with the lowest ORF1ab Ct values and a genome coverage >96% were selected from each of B.1.1.7 and B.1.177 (EU1) lineage for use in this work.

### RT-PCR

Reverse transcription was carried out using the SuperScript VILO cDNA synthesis kit (Invitrogen) at 25 ºC for 10 minutes, then 50 ºC for 10 minutes, followed by 2 minutes each at 55 ºC, 50 ºC, 60 ºC, 50 C, 65 ºC, and 50 ºC. PCR was carried out using GoTaq DNA polymerase (Promega) with RT-PCR primers 5′UTR-FWD and ORF9-REV (see “Primer Sequences”, below). PCR cycling conditions were 95 ºC for 2 minutes, followed by 40 cycles of 95 ºC for 30 s, 56 ºC for 30 s, and 72 ºC for 60 s, with a final extension of 72 ºC for 5 minutes. PCR products were purified using a NucleoSpin Gel and PCR Clean-up kit (Macherey-Nagel). PCR products were separated in 1.2% agarose-TBE gels at 110 V for 45 minutes, before visualisation using an Amersham Imager 600.

### Nanopore Sequencing and Data Analysis

End preparation was carried out using the NEBNext Ultra II End Repair A-Tailing Module (NEB). The reaction was carried out at 20 ºC for 25 minutes, followed by 15 minutes at 65 ºC to inactivate the enzyme. Samples were directly ligated with an individual barcode from the ONT Native Barcoding kit (EXP-NBD196) using NEBNext Blunt/TA Ligase Master Mix (NEB). The ligation reaction was carried out for 20 minutes at 20 ºC followed by 10 minutes at 70 ºC. After barcoding, samples were pooled with equal volumes. The pool is cleaned up using a 0.4x ratio of SPRIselect magnetic beads (Beckman Coulter). Fragment-bound beads were washed twice with Short fragment buffer (SFB) (ONT), and once with 80% ethanol. The DNA target was eluted with Buffer EB (Qiagen). To allow Nanopore sequencing, the AMII adapter (ONT) was ligated on using NEBNext Quick Ligation Module (NEB), with a 20 minute incubation at 20 ºC. A second SPRIselect bead cleanup was carried out with a 1x ratio of beads and two washes with SFB only. The final pool was eluted in Elution Buffer (ONT) and quantified by Qubit HS dsDNA assay (Thermo Fisher). A FLO-MIN106 flowcell is primed with the Flow cell Priming kit (ONT). Up to 15ng of the final pool was mixed with Loading Beads (ONT) and Sequencing Buffer (ONT) before loading onto the flowcell, which was sequenced on the GridION platform for up to 20 hours with a voltage of −180 mV. Sequencing reads were adapter-trimmed using porechop^37^ and reads were filtered by size into bins of 75-130bp, 250-400bp, and 850-1050bp, and aligned to the reference sequence of the ORF9 amplicon using minimap2^38^ and samtools^39^. Alignments were visualised in Integrative Genomics Viewer^40^, and statistics were extracted using the samtools flagstat command. ORF9b-specific amplicons in were identified by counting the number of reads in the 850-1050 bin containing the ORF9b sgmRNA-specific sequence TGTAGATCTGTTCTCTAAATGGACC. A consensus sequence was generated from reads in 75-130bp bin, which identified this product as a result of mis-priming of 4 bases at the 3′-end of the reverse PCR primer. Quantification of read statistics were plotted using GraphPad Prism.

### RT-qPCR

To quantify sgmRNA abundance, RT-qPCR was carried out using the Taqman Multiplex Master Mix (Applied biosystems) with 1.8 µM forward and reverse primers and µM 5′-FAM-/3′-BHQ-labelled probe and cDNA from 5 ng of RNA extracted from infected VeroE6 cells (or a 1:100 dilution for N), or from 2 µL of RNA from HCW swab samples. Linear amplification range was tested against synthetic oligonucleotide templates (Dharmacon) and absolute sgmRNA copy number was calculated by interpolating cDNA standard curves. Ratios between sgmRNA species were calculated from copy numbers. ORF1ab abundance was determined using the Real-Time Fluorescent RT-PCR kit for Detecting nCoV-19 (BGI). Log10-transformed data from infected cells were compared to B Lineage-infected cells by one-way ANOVA with Dunnett’s test for multiple comparisons using GraphPad Prism (v 9.2.0). To account for differences in variances due to uneven samples sizes from swab samples, data were compared to EU1 swabs by one-way Browne-Forsythe and Welch ANOVA with Dunnett’s T3 test. To determine innate immune responses, RT-qPCR was carried out using the Brilliant III Ultra-Fast SYBR Green QPCR Master Mix (Agilent) with 0.8 µM primers, against IFNb, IFIT1 or GAPDH (see “Primer Sequences”, below), and cDNA from 5 ng of RNA extracted from transfected HEK293T cells. Data were normalised to GAPDH and expressed as fold change (2-DDCq) over control cells, which were not transfected with poly(I:C), for each plasmid. Log10-transformed data were compared by one-way ANOVA with Dunnett’s test for multiple comparisons using GraphPad Prism (v 9.2.0).

### Immunoblotting

Cell lysates (6-10 µg total protein) were separated by SDS-PAGE using Any kD precast gels (Bio-Rad) and transferred to 0.2-µm nitrocellulose membrane by semidry electrotransfer. Membranes were probed with anti-Nucleocapsid (MA5-35943, Invitrogen, 1:400), anti-HSP90 (MA5-35624, Invitrogen, 1:2000) or anti-Membrane (MRC PPU & CVR Coronavirus Toolkit, Sheep No. DA107, 1:800), followed by IRdye secondary antibodies (Li-Cor), allowing visualisation on an Odyssey CLx imaging system (Li-Cor).

### Transfection

HEK293T cells were transfected at 70 % confluency in 24-well plates, with 25, 50 or 100 fmol pCDNA3-B117-N-T2A-mCherry or pCDNA3-B117-Nstar-T2A-mCherry from B.1.1.7, or 100 fmol pCDNA-NS1, or mock transfected, using Lipofectamine 2000 (Invitrogen) at a 2:1 ratio according to the manufacturer’s instructions, in antibiotic-free DMEM. After 24 hours, cells were transfected again with 1 µg Polyinosinic–polycytidylic acid (Poly(I:C), Sigma Aldrich) using Lipofectamine 2000 at a 2:1 ratio in Opti-MEM (Gibco). After a further 24 hours, cells were harvested in passive lysis buffer for western blot analysis (Promega) and RNA was extracted by mixing cell lysate 1:3 with TRIzol LS (Invitrogen) then using a Direct-zol RNA MiniPrep kit (Zymo Research).

### Phylogenetics and identification of novel TRS-B sites

For Fig. 1a and Extended Data Fig. 3a phylogenetic trees produced by Nextstrain for SARS-CoV-2 global data were downloaded as SVG files under the CC BY license on 23/09/2021, following annotation to indicate amino acids at position 204 of N, using the Colour by… genotype command. A complete snapshot of the GISAID EpiCov database was downloaded on 12 Nov 2021 and the subset of ‘complete’, ‘high coverage’ SARS-CoV-2 genomes extracted according to the associated metadata. These were aligned in parallel in batches of 10000 sequences to the Wuhan-Hu-1 (NC_045512) reference with MAFFT^41^ (‘mafft –nuc –nwildcard – 6merpair --keeplength –addfragments’) and the results merged to produce a complete alignment. Locations of the AAACGAAC and ACGAAC sequence motifs were extracted from individual sequences within the alignment using ‘seqkit locate’ ^42^ using regular expressions allowing for alignment gaps (‘-’ characters) within each search sequence. These were subsequently compared to the locations of the same search sequences within the Wuhan-Hu-1 reference using ‘bedtools subtract’ to identify novel occurrences and their frequencies calculated with ‘bedtools genomecov’. As the ‘--keeplength’ flag to MAFFT carries the potential to create artefactual alignments by removing nucleotides inserted relative to the reference, for each novel location of either sequence motif, the corresponding genomes were extracted and re-aligned to the Wuhan-Hu-1 reference using MAFFT without this flag to ensure that reported sites were valid.

To screen for possible convergent evolution, we examined the Audacity tree made available by the GISAID Initiative^34^, using a mutation-annotated tree inferred by UShER^43^. We examined the final phylogenetic tree using Taxonium (https://github.com/theosanderson/taxonium), searching for nodes annotated with the mutations under consideration, and which had more than 30, 50, or 100 descendants. Where clades were found, we considered the alternative possibilities of convergent evolution, phylogenetic misplacement, or artefacts due to contamination with other B.1.1 sequences. In particular we looked for the presence of mutations back to reference, often indicative of artifacts due to contamination or failure to trim primer sequences, and used CoV-Spectrum^44^ to examine sub-clade dynamics and prevalence.

## Primer Sequences

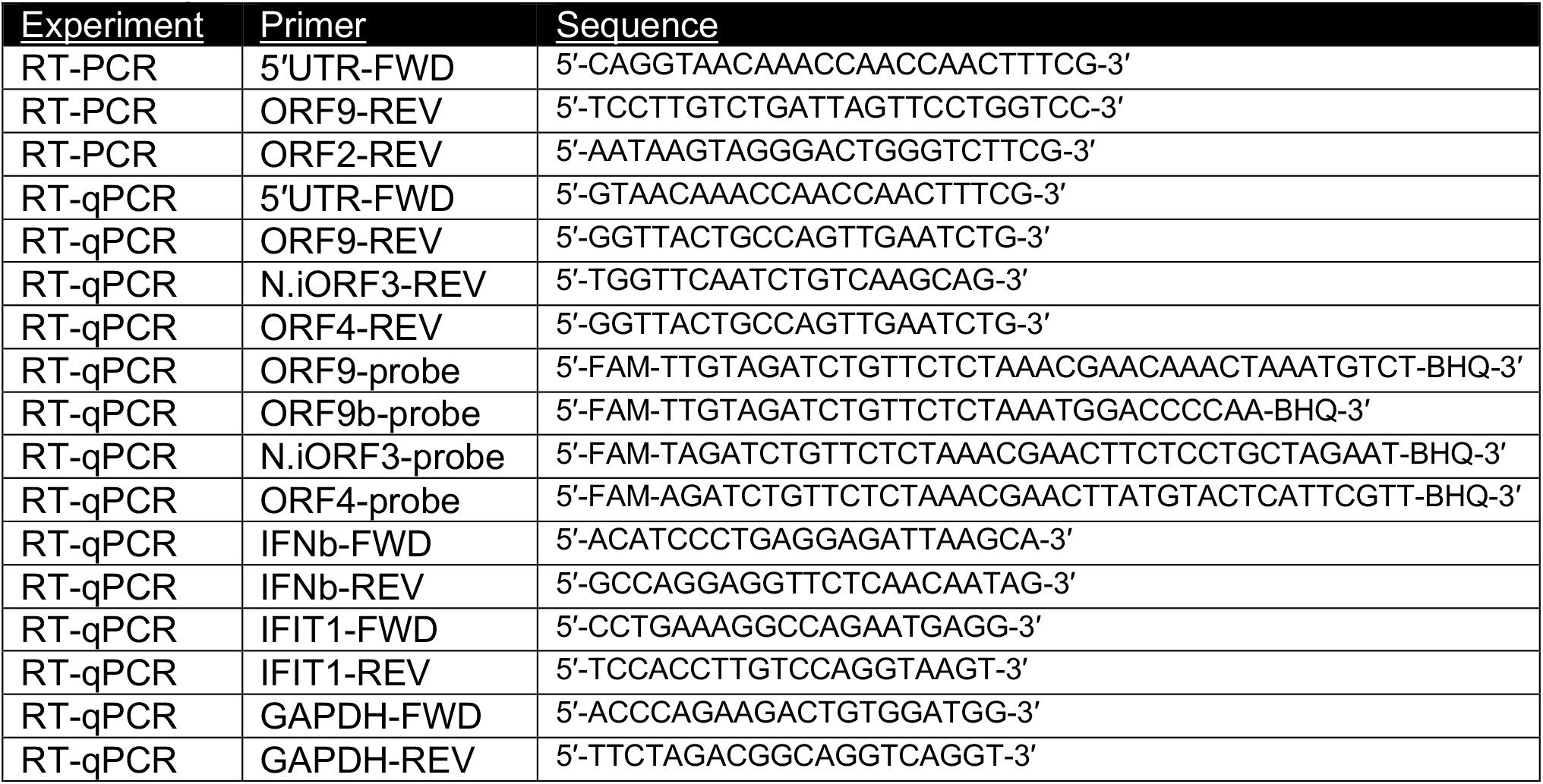

---

**Extended Data Fig. 1.**
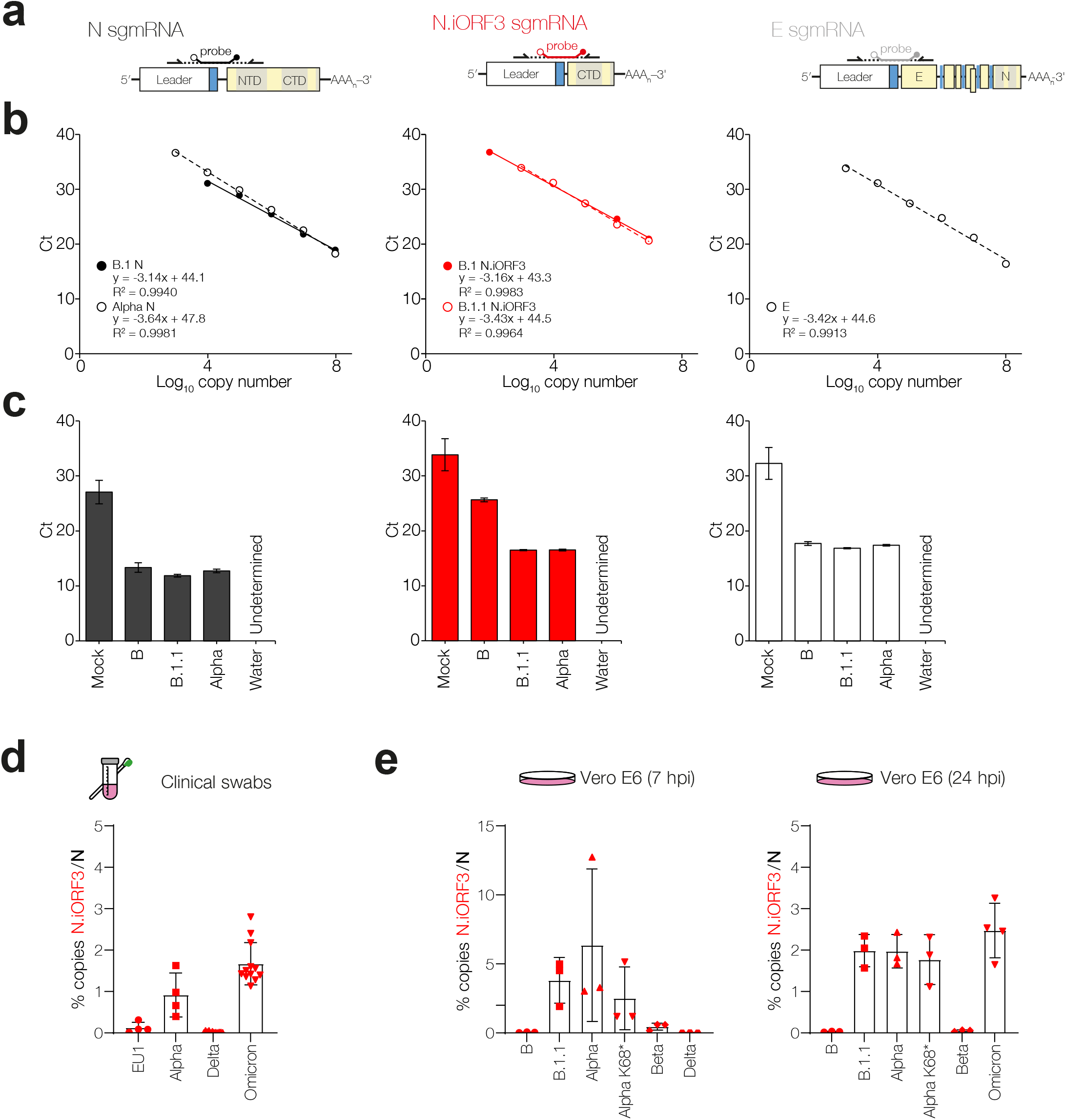
Validation of sgmRNA RT-qPCR and determination of sgmRNA copy number. **a**, Schematic representation of qRT-PCR primer probe sets for N, N.iORF3 and E sgmRNAs, and **b**, standard curves using synthetic cDNA oligonucleotide templates. Insets show linear regression of Ct plotted against log_10_-transformed cDNA copy number. **c**, RT-qPCR of VeroE6 cells infected with B, B.1.1, B.1.1.7 or mock cells, or water controls, validating RT-qPCR specificity. Bars represent the mean and standard deviation of three biological replicates. RT-qPCR analysis of N.iORF3 sgmRNA copy number, expressed as a ratio of N copy number in **d**, clinical swabs, and **e**, infected VeroE6 cells in culture at 7 and 24 hours post-infection. For infected VeroE6 cells, data are means and standard deviations of at least three biological replicates. For clinical swabs, data are means and standard deviations of four (EU1/Alpha) or twelve (Delta/Omicron) swab samples per lineage.

**Extended Data Fig. 2.**
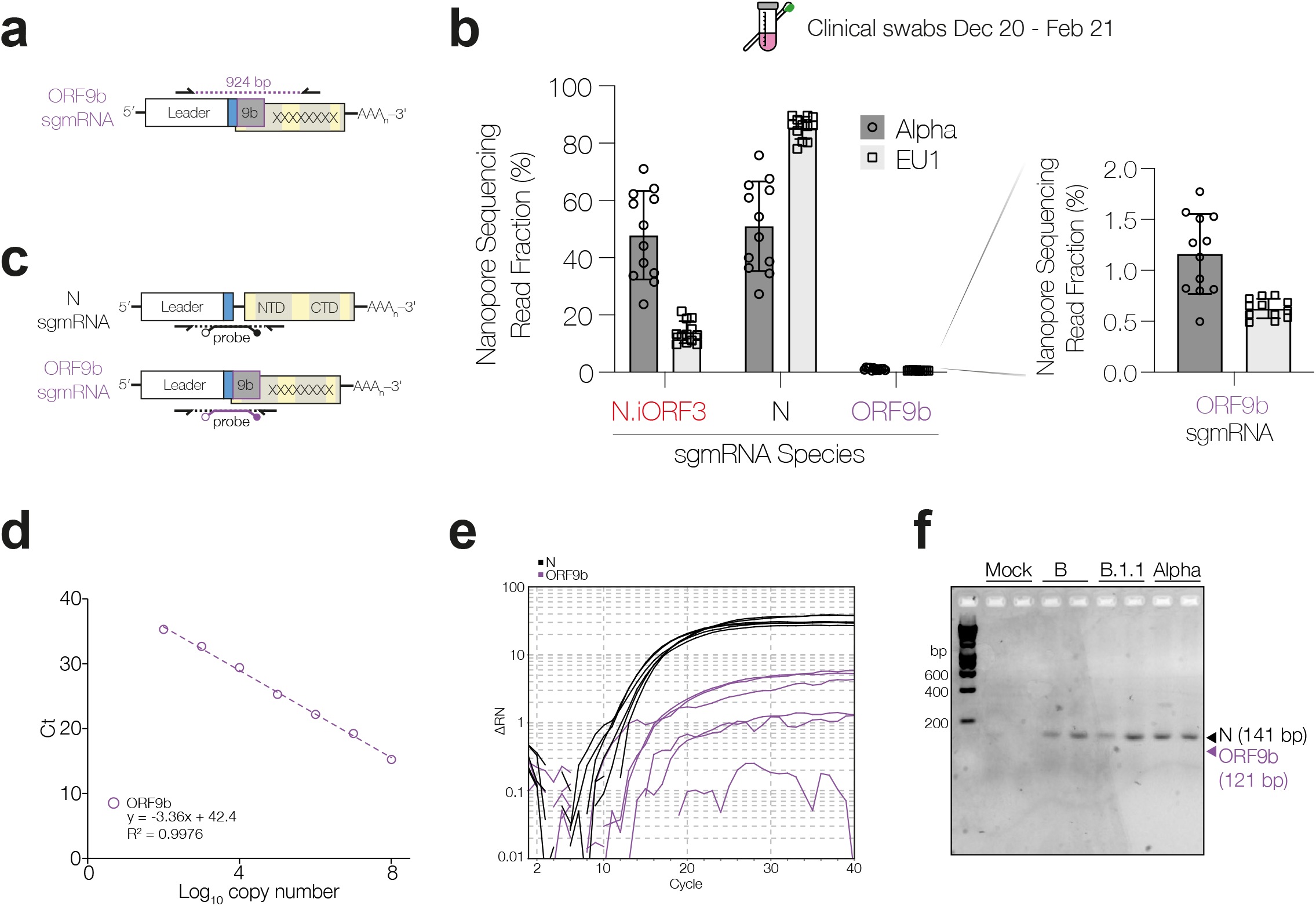
Detection of ORF9b-specific sgmRNA in infected cells and human swab samples. **a**, Schematic for detection of ORF9b-specfic sgmRNA using the endpoint RT-PCR assay depicted in Fig. 2a. **b**, Fraction of endpoint PCR products from N.iORF3-, N-, or ORF9b-specific sgmRNAs in clinical swab samples from EU1 and Alpha lineages, as determined by nanopore sequencing. Data are means and standard deviations of twelve swab samples per lineage. **c**, Schematic representation of qRT-PCR primer probe sets for N and ORF9b sgmRNAs, and **d**, standard curve using synthetic cDNA oligonucleotide templates for ORF9b sgmRNA. Standard curve for N sgmRNA is shown in Extended Data Fig. 1b. **e**, Amplification of N (black) or ORF9b (purple) sgmRNA in Vero E6 cells infected with B, B.1.1 or B.1.1.7, in technical duplicate, representative of three biological replicates. **e**, Gel electrophoresis of RT-PCR products from ORF9b RT-qPCR reactions.

**Extended Data Fig. 3.**
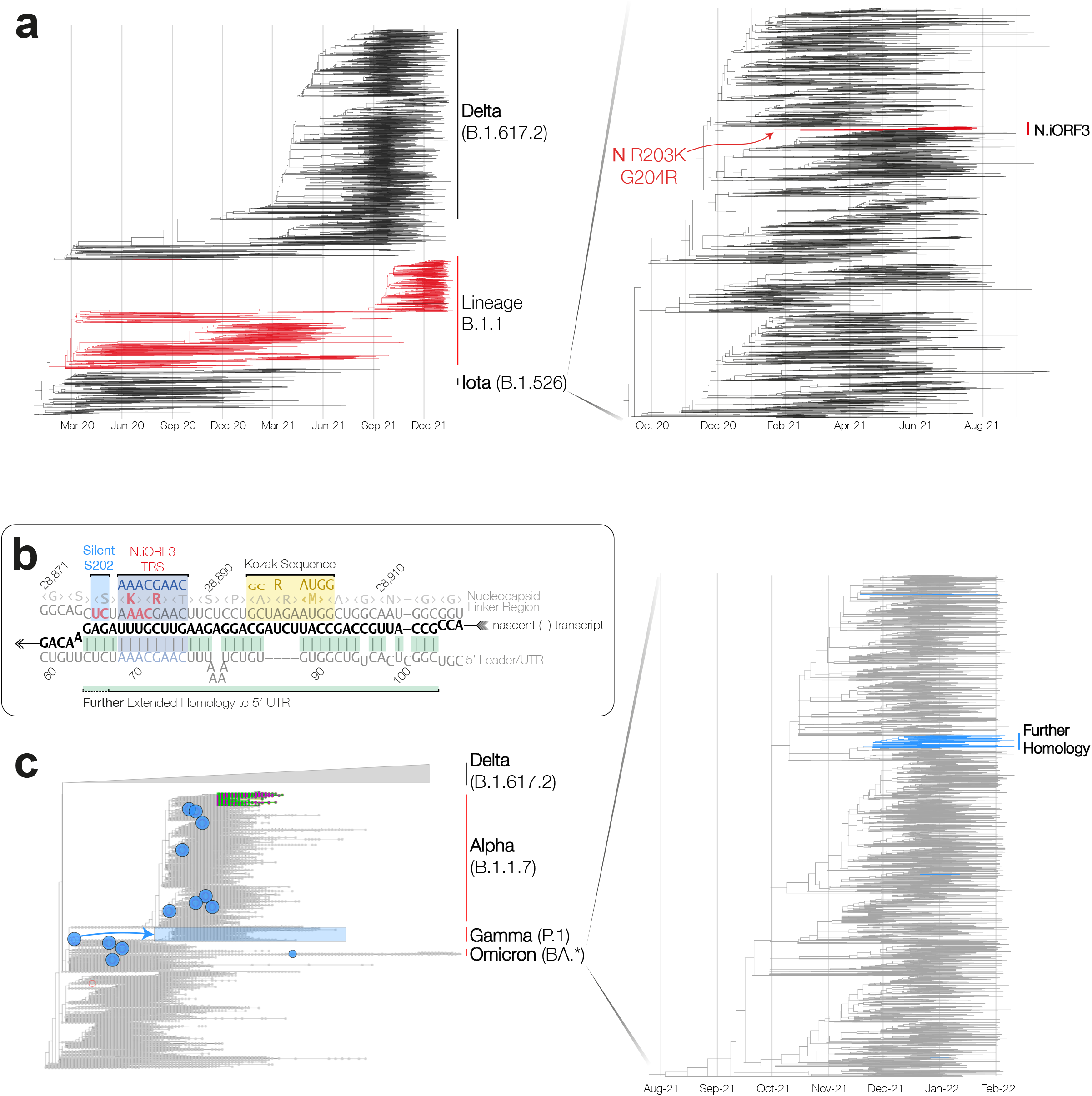
N.iORF3 and extended homology to TRS-flanking regions have both evolved convergently. **a**, Phylogenetic reconstruction of the Iota variant evolution (B.1.526), highlighting the emergence of N.iORF3 (red). **b**, A silent mutation at S202 generates extended homology of the N.iORF3 TRS-B region to the 5’UTR region flanking the TRS-L. **c**, Phylogenetic reconstruction of SARS-CoV-2 evolution in humans, with independent emergences of extended N.iORF3 TRS sequence highlighted (See **Extended Data Table 1**). The phylogenetic tree in panel a was adapted from Nextstrain based on the Iota-focused build, and in panel c on the Omicron.21K focused-build, both maintained by Emma Hodcroft and Richard Neher^32,33^.

**Extended Data Fig. 4.**
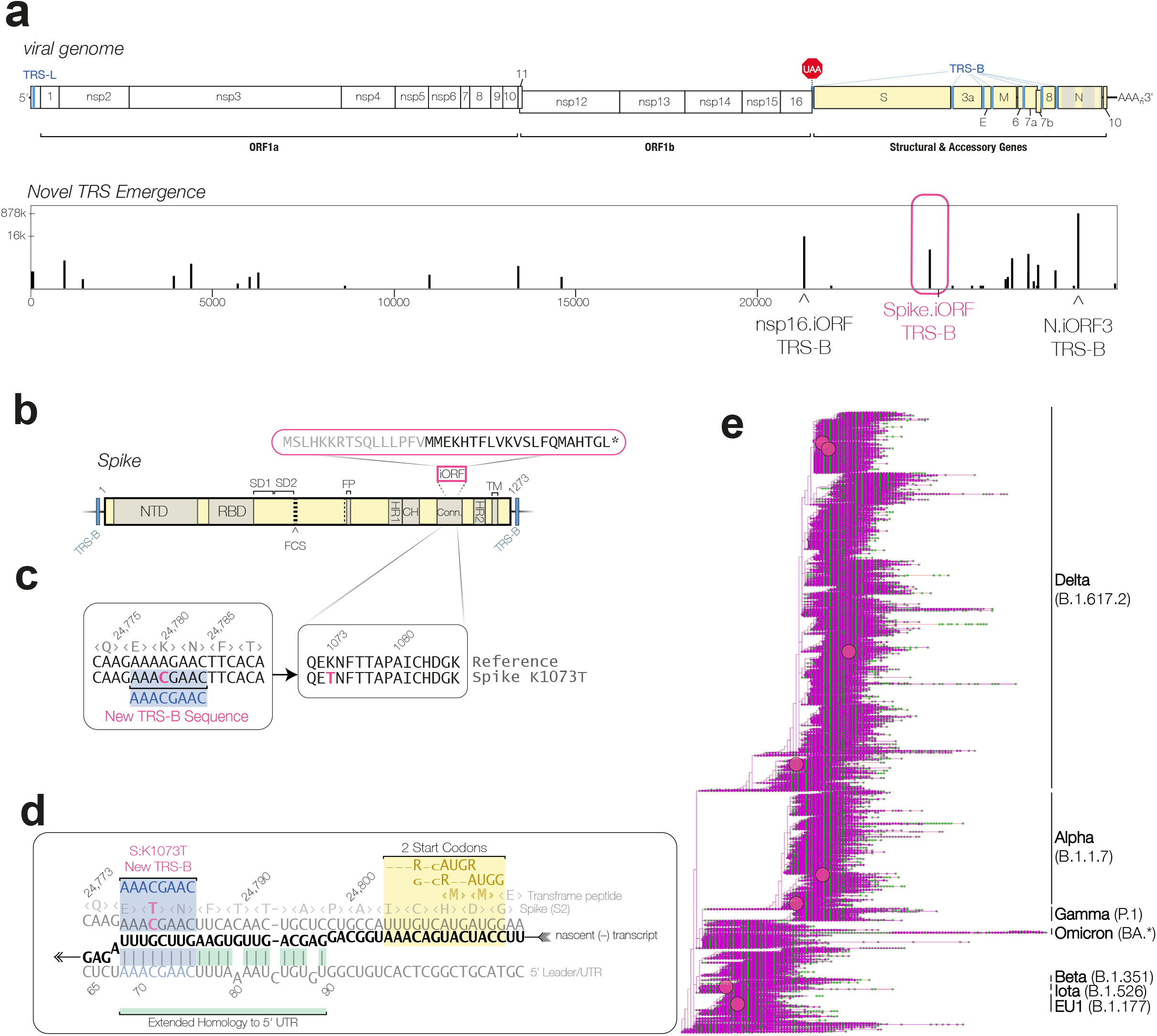
Convergent evolution of a TRS-B site within the coding region of Spike protein, overlapping the connector domain (CD) of Spike/S2. **a**, Schematic of the SARS-CoV-2 genome (upper panel) and frequency of emergence of the TRS-B sequence (AAACGAAC) in the global SARS-CoV-2 population (lower panel). **b**, Diagram of the Spike ORF, including a potential transframe product. c, Sequence alignment of amino acids 1071-1086 of Spike, and alignment of the corresponding nucleotide sequences show emergence of a new TRS-B sequence. **d**, The sequence context of the novel Spike.iORF sgmRNA, showing TRS-B (blue highlight), extended homology to the 5’UTR (green highlight) during nascent (-) strand RNA synthesis (black), and downstream tandem start codons and Kozak contexts (yellow highlight). **e**, Phylogenetic reconstruction of SARS-CoV-2 evolution in humans, with independent emergences of Spike.iORF TRS sequence with 250 descendant genomes highlighted in pink (See **Extended Data Table 1**). The schematic and abbreviations of Spike protein domains shown in panel b was adapted from Lan et al.^16^

**Extended Data Fig. 5.**
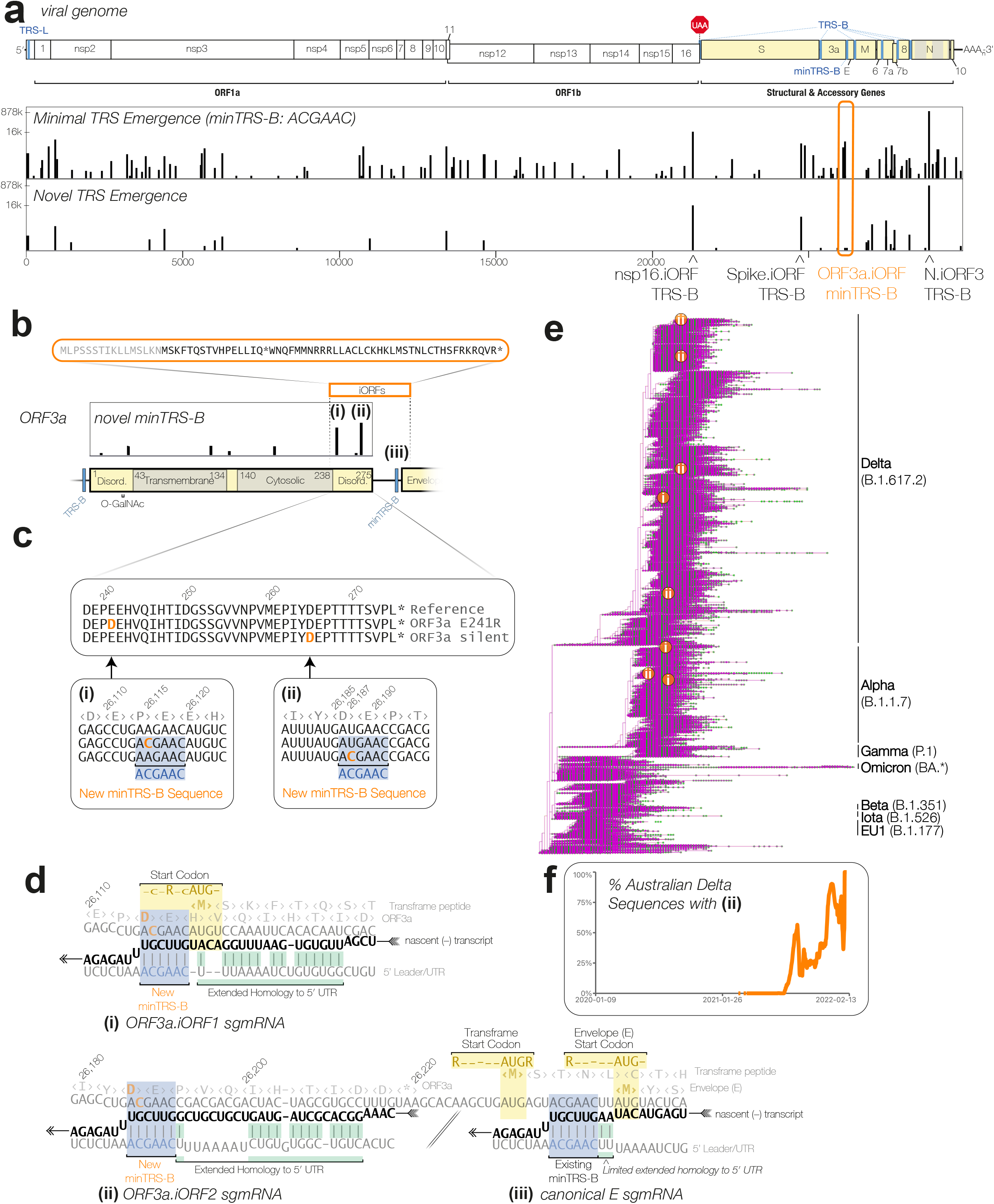
Minimal TRS-B (minTRS-B) sites have evolved independently within ORF3a, upstream of the canonical minTRS-B driving Envelope (E) sgmRNA expression. **a**, Schematic of the SARS-CoV-2 genome (upper panel) and frequency of emergence of the minTRS-B sequence (ACGAAC, middle panel) and full-length TRS-B sequence (AAACGAAC, lower panel) in the global SARS-CoV-2 population. **b**, Diagram of the ORF3a, including a potential transframe product, showing two loci of new minTRS-B emergence (i) and (ii), upstream of (iii) the canonical E minTRS-B. **c**, Sequence alignment of amino acids 238-275 of ORF3a, and alignment of the corresponding nucleotide sequences show two loci of new minTRS-B emergence. **d**, The sequence context of the novel (i) ORF3a.iORF1 and (ii) ORF3a.iORF2 sgmRNA relative to (iii) existing minTRS-B that drives canonical E sgmRNA expression, showing minTRS-B sites (blue highlights), extended homologies to the 5’UTR (green highlights) during nascent (-) strand RNA synthesis (black), and downstream start codons and Kozak contexts (yellow highlights). **e**, Phylogenetic reconstruction of SARS-CoV-2 evolution in humans, with independent emergences of ORF3a.iORF minTRS-B sites with 250 descendant genomes highlighted in orange, with overlaid (i) or (ii) annotation in white denoting ORF3a.iORF1 minTRS-B and ORF3a.iORF2 minTRS-B emergence, respectively. **f**, Percentage of sequenced Delta variant genomes in Australia that contain (ii) ORF3a.iORF2 minTRS-B mutation (See **Extended Data Table 1**). The plot in panel f was generated using CoV-Spectrum^44^.

**Extended Data Table 1.**
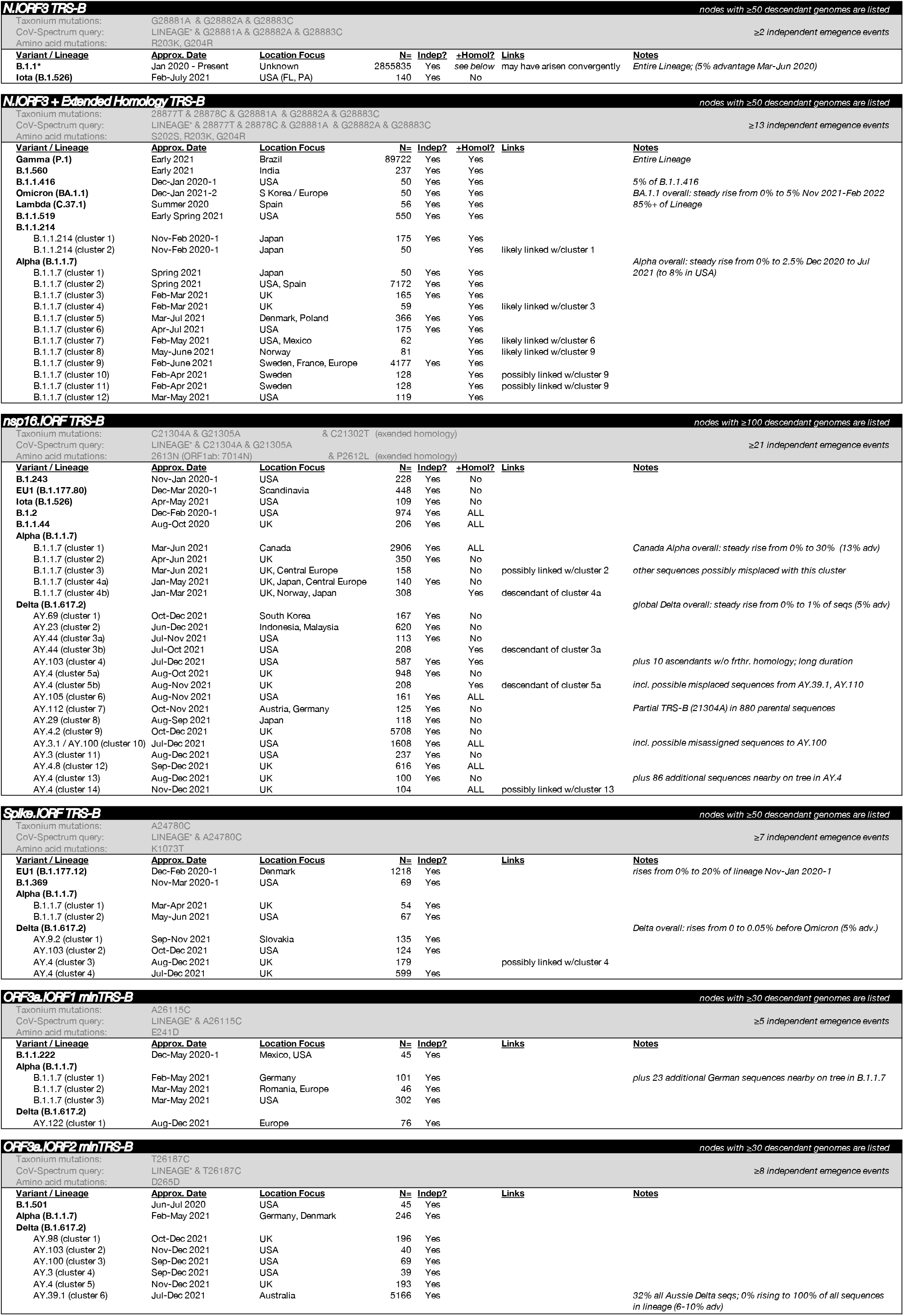
Convergent evolution of TRS-B and minTRS-B sites within SARS-CoV-2. Identified nodes on a phylogenetic reconstruction of SARS-CoV-2 evolution in humans representing clusters of novel TRS and sgmRNA emergence (see Methods). For each emergence event, the parental lineage, approximate date (“Approx. Date”) and location (“Location Focus”) of circulation are annotated, as well as number of identified descendant genomes within each cluster (“N=“), whether it appears to be a true independent emergence event (“Indep?”, see Methods) and whether it has further extended homology to the 5’UTR (“+Homol?”). Each cluster is further annotated with potential interrelationships (“Links”) and additional notes, including context to its spread based on analysis using CoV-Spectrum^44^.

